# scRNA-sequencing reveals subtype-specific transcriptomic perturbations in DRG neurons of *Pirt-EGFPf* mice in neuropathic pain condition

**DOI:** 10.1101/2022.01.06.475187

**Authors:** Chi Zhang, Ming-Wen Hu, Shao-Qiu He, Xue-Wei Wang, Xu Cao, Feng-Quan Zhou, Jiang Qian, Yun Guan

## Abstract

Functionally distinct subtypes/clusters of dorsal root ganglion (DRG) neurons, which differ in soma size and neurochemical properties, may play different roles in nerve regeneration and pain. However, details about transcriptomic changes in different neuronal subtypes under maladaptive neuropathic pain conditions remain unclear. Chronic constriction injury (CCI) of the sciatic nerve represents a well-established model of neuropathic pain that mimics the etiology of clinical conditions. Therefore, we conducted single-cell RNA-sequencing (scRNA-seq) to characterize subtype-specific perturbations of transcriptomes in lumbar DRG neurons 7 days after sciatic CCI. By using *Pirt*-*EGFPf* mice that selectively express enhanced green fluorescent protein in DRG neurons, we established a highly efficient purification process to enrich neurons for scRNA-seq. We observed a loss of marker genes in injured neurons of 12 standard neuronal clusters, and the emergence of four prominent CCI-induced clusters at this peak-maintenance phase of neuropathic pain. Importantly, a portion of injured neurons from a subset of the 12 standard clusters (NP1, PEP5, NF1, and NF2) were spared from injury-induced identity loss, suggesting subtype-specific transcriptomic changes in injured neurons. Moreover, uninjured neurons, which are necessary for mediating the evoked pain, also demonstrated subtype-specific transcriptomic perturbations in these clusters, but not others. Notably, male and female mice showed differential transcriptomic changes in multiple neuronal clusters after CCI, suggesting transcriptomic sexual dimorphism in primary sensory neurons after nerve injury. Collectively, these findings may contribute to the identification of new target genes and development of DRG neuron subtype-specific therapies for optimizing neuropathic pain treatment and nerve regeneration.

## Introduction

The dorsal root ganglion (DRG) contains the somas of primary sensory neurons, which differ in size and axon myelination. Different gene expression profiles confer divergent neurochemical, physiologic, and functional properties on the various subtypes of DRG neurons (Gatto et al., 2019; Sharma et al., 2020; Usoskin et al., 2015; Zeisel et al., 2018; Zheng et al., 2019). Small-diameter neurons are important for transmitting nociceptive and thermal information, whereas large-diameter neurons are mainly non-nociceptive neurons, including mechanoreceptors and proprioceptors. Nerve injury induces various responses in DRG neurons, including cell stress, regeneration, hyperexcitability, and functional maladaptation. How these changes vary in functionally distinct neuronal subtypes and possibly affect nerve regeneration and neuropathic pain remains unclear.

Recently, single-cell/single-nucleus RNA-sequencing (scRNA-seq/snRNA-seq) has begun to reveal transcriptomic perturbations in DRG neurons after transection or crush nerve injury (Hu et al., 2016; Renthal et al., 2020). Although these models are suitable for studying nerve regeneration, they may not closely capture the etiology of clinical neuropathic pain, which often involves chronic compression, neuroinflammation, and partial injury to a major nerve. Chronic constriction injury (CCI) of the sciatic nerve represents a well-established neuropathic pain model that encompasses compression, ischemia, inflammation, and axonal demyelination (Bennett & Xie, 1988). It mimics the etiology of clinical conditions and induces symptoms similar to those of post-traumatic neuropathic pain in humans (e.g., tactile allodynia, spontaneous pain) (Challa, 2015). Accordingly, sciatic CCI represents one of the most generalizable neuropathic pain models (Costigan et al., 2010; Griffin et al., 2007; LaCroix-Fralish et al., 2011).

Axotomized neurons may exhibit the most profound gene expression changes that are important for regeneration. Nevertheless, neighboring uninjured DRG neurons also show significant functional changes (e.g., hyperexcitability) and contribute to dysesthesia and evoked pain hypersensitivity as a result of the remaining peripheral innervations (Djouhri et al., 2012; Kalpachidou et al., 2021; Obata et al., 2003; Tran & Crawford, 2020). Details of subtype-specific changes in DRG neurons under neuropathic pain conditions are currently only partially known. In particular, it is unclear how nerve injury affects transcriptomes in uninjured neurons at the single-cell level. Moreover, increasing clinical and preclinical evidence suggests that males and females have differences in pain sensitivity and susceptibility to chronic pain. To optimize clinical treatment, it will be important to delineate sex-related gene expression changes in functionally distinct subtypes of DRG neurons after nerve injury, and determine how these changes underpin sexual dimorphisms in neuropathic pain.

We established a highly efficient purification approach by using *Pirt*-*EGFPf* mice to enrich DRG neurons for scRNA-seq. *Pirt* is expressed in >83.9% of neurons in mouse DRG, but not in other cell types (Kim et al., 2008). Thus, green fluorescent protein (GFP) is selectively expressed in most DRG neurons in *Pirt*-*EGFPf* mice, driven by the *Pirt* promotor. Thus, GFP expression allows effective purification of DRG neurons from these mice. We then characterized perturbations of transcriptomes in DRG neurons after sciatic CCI. Because only a portion of nerve fibers are injured after CCI, the lumbar DRGs contain a mixture of injured and uninjured neurons. In addition to characterizing injured neurons, which express high levels of the injury marker gene *Sprr1a*, we also examined subtype-specific transcriptomic changes in uninjured neurons (*Sprr1a-*) and transcriptomic sexual dimorphism after nerve injury. Our findings may be useful for developing DRG neuron subtype-specific treatment and sex-specific therapies for nerve regeneration and neuropathic pain.

## Materials and Methods

### *Pirt*-*EGFPf* mouse

The *Pirt*-*EGFPf* mouse strain was a gift from Dr. Xinzhong Dong in the Solomon H. Snyder Department of Neuroscience, School of Medicine, Johns Hopkins University (Baltimore, MD). Adult mice (7-8 weeks old) of both sexes were used. Genotypes of the mice were determined by PCR using primers provided by the Mutant Mouse Resource & Research Centers. Mice were housed 3-5 per cage and given access ad libitum to food and water. All animal experiments were conducted in accordance with the protocol approved by the Institutional Animal Care and Use Committee of the Johns Hopkins University.

### Bilateral Sciatic Nerve CCI

Contralateral changes may develop after unilateral CCI of the sciatic nerve, including spontaneous pain and mechanical hypersensitivity in hind paws (Paulson et al., 2002; Wilkerson et al., 2020), as well as gene expression in the spinal cord and DRGs (Jancalek et al., 2010). Because transcriptional changes in contralateral DRGs may differ from those on the ipsilateral side, we performed bilateral sciatic CCI to allow pooling of bilateral DRG tissues for sequencing and to avoid sample variations. The bilateral CCI model has been validated in previous studies, which showed that animals displayed prolonged cold and mechanical hypersensitivity in hind paws but did not exhibit the asymmetric postural or motor influences of unilateral CCI or behavior changes (Dai et al., 2014; Datta et al., 2010; Vierck et al., 2005).

Adult *Pirt*-*EGFPf* mice were randomly assigned to undergo bilateral CCI surgery or sham surgery. All procedures were performed by the same experimenter to avoid variation in technique. CCI of sciatic nerve was performed as previously described (Guan et al., 2010; Li et al., 2017). Briefly, mice were anesthetized with 2% isoflurane, and a small incision was made at mid-thigh level. The sciatic nerve was exposed by blunt dissection through the biceps femoris. The nerve trunk proximal to the distal branching point was loosely ligated with three nylon sutures (9-0 nonabsorbable monofilament; S&T AG) placed approximately 0.5 mm apart until the epineuria was slightly compressed and hindlimb muscles showed minor twitching. The muscle layer was closed with 4-0 silk suture and the wound closed with metal clips. *Pirt-EGFPf* mice developed mechanical hypersensitivity in both hind paws, as indicated by a significant increase in paw withdrawal frequency (PWF) to von Frey filament stimulation (**Figure 1B**).

**Figure 1.**
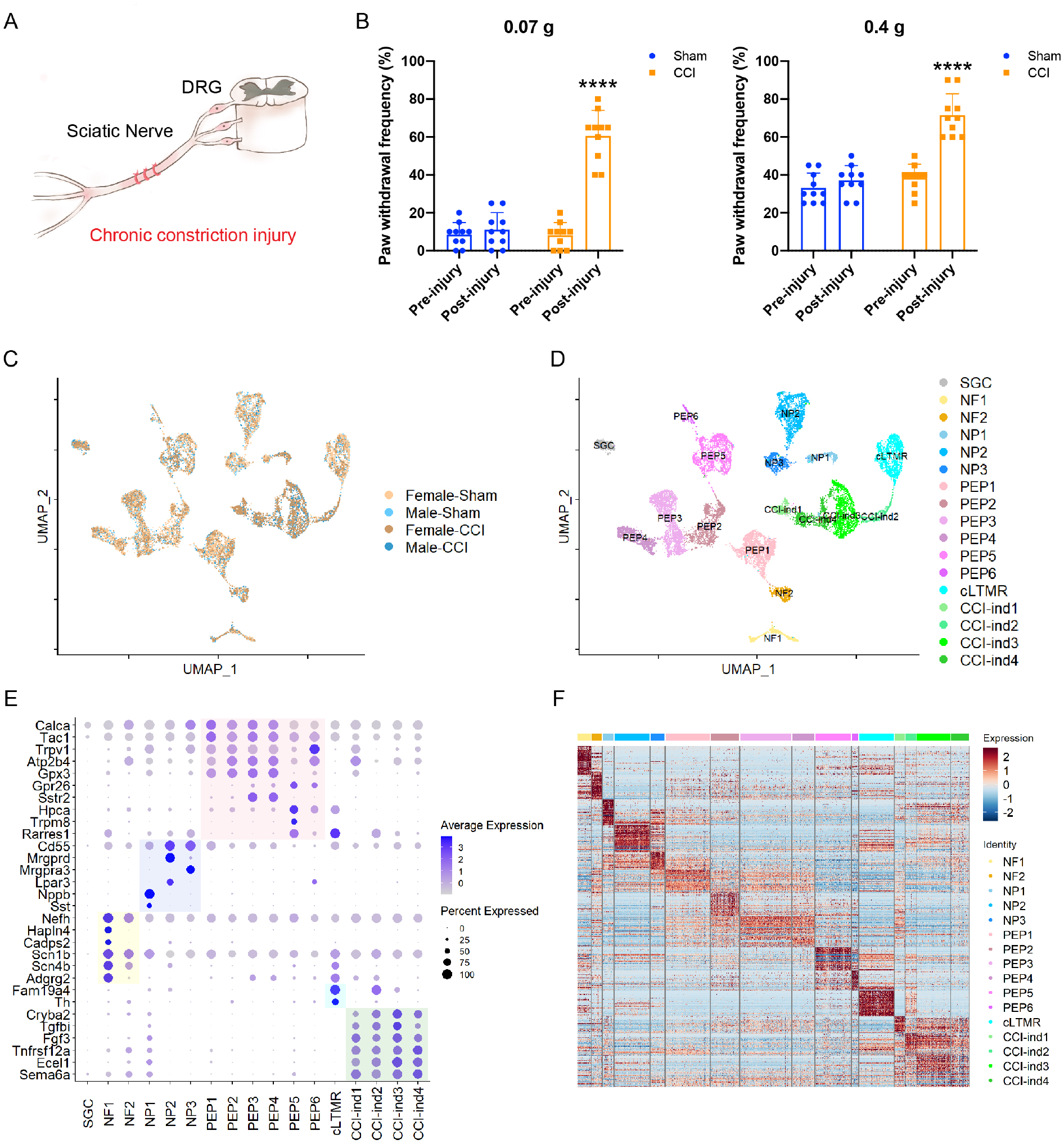
scRNA-seq identified distinct clusters of cells in the dorsal root ganglion (DRG) of *Pirt*-*EGFPf* mice. (A) Schematic diagram showing procedure for CCI of the sciatic nerve. (B) Paw withdrawal frequencies to low-force (0.07 g von Frey filament, left) and high-force (0.4 g, right) mechanical stimuli before and 6 days after CCI or sham surgery. \*\*\*\**P* < 0.0001 vs pre-injury. Two-way mixed-model ANOVA (Bonferroni post hoc test). Data are expressed as mean + SD. (C) Integration of four datasets visualized by UMAP. (D) Seventeen distinct cell clusters were identified by Seurat, including SGC (1), NF (2), NP (3), PEP (6), cLTMR (1), and CCI-induced clusters (4). (E) Dot plot of subtype-specific marker genes in each cluster. Genes highlighted in the yellow, purple, pink, and blue zones are known markers for NF, NP, PEP, and cLTMR, respectively. Genes highlighted in the green zone are markers identified in CCI-induced clusters. The dot size represents the percentage of cells expressing the gene, and color scale indicates the average normalized expression level in each cluster. (F) A heatmap shows the expression patterns of the top 50 marker genes in each cluster.

### Mechanical Hypersensitivity Test

Animals were allowed to acclimate for a minimum of 48 h before any experimental procedures. Hypersensitivity to punctuate mechanical stimuli was assessed by the PWF method using 2 von Frey monofilaments (low-force, 0.07 g; high-force, 0.4 g). Each von Frey filament was applied perpendicularly to the midplantar area of each hind paw for ~1 s. The left hind paw was stimulated first, followed by the right side (>5-min interval). The stimulation was repeated 10 times at a rate of 0.5 to 1 Hz (1- to 2-s intervals). If the animal showed a withdrawal response, the next stimulus was applied after the animal resettled. PWF was then calculated as (number of paw withdrawals/10 trials) × 100%.

### Single cell dissociation

Bilateral L4-5 DRGs were collected from mice at day 7 after bilateral sciatic CCI or sham surgery. Male and female mice from the same litter were subjected to the same surgery (sham or CCI). DRGs from the same group of mice were pooled and defined as one sample for sequencing. The four groups included Female-Sham, Male-Sham, Female-CCI, and Male-CCI. DRGs were dissected out, digested with 1 mg/ml type I collagenase (Thermo Fisher Scientific) and 5 mg/ml dispase II (Thermo Fisher Scientific) at 37°C for 70 min (10 DRGs/tube), and disassociated into single cells in Neurobasal medium containing 1% bovine serum albumin (BSA). Cells were filtered through a 40 μm cell strainer and centrifuged at 500×g for 5 min. The pellet was resuspended in 500 μl of Triple Express (Thermo Fisher Scientific) and digested at 37°C for 2 min. The reaction was stopped with 1 ml of fetal bovine serum. The cell suspension was laid onto 5 ml of Neurobasal medium containing 20% Percoll and centrifuged at 1,400 RPM for 8 min. The upper layer was removed first and then the lower layer. The pellet was washed with 2 ml of Neurobasal medium containing 1% BSA and centrifuged at 500×g for 5 min. The pellet was resuspended with 500 μl of Neurobasal medium containing 1% BSA. The GFP^+^ cells were sorted into 500 μl of Neurobasal medium containing 1% BSA and centrifuged at 500×g for 5 min to remove most supernatant. The pellet was resuspended with the remaining supernatant to a concentration of ~1000 cells/μl.

### 10x Genomics Library Preparation and Sequencing

The single cell suspensions were further processed with Chromium Next GEM Single Cell 3′ GEM, Library & Gel Bead Kit v3 (PN-1000094) according to the manufacturer’s instructions to construct the scRNA-seq library. All libraries were sequenced with the Illumina NovaSeq platform. The raw sequencing reads were processed by Cell Ranger (v.2.1.0) with the default parameters. The reference genome was mm10.

### Single-cell RNA-seq Data Analysis

Scrublet with default parameters was used first to remove single-cell doublets. After doublet removal, we filtered out cells with fewer than 1,500 genes expressed and cells with more than 10% mitochondrial UMI counts. The rigorous filtering was intended to remove smaller residual non-neuronal cells such as SGCs. After doublet removal and quality control, we applied Seurat’s integration workflow to correct possible batch effect for the remaining cells of the four datasets. Prior to the integration, the four datasets were transformed into four individual Seurat objects with standard steps including “CreateSeuratObject,” “NormalizeData,” and “FindVariableFeatures.” Subsequently, we used “FindIntegrationAnchors” with the top 3,000 variable genes to locate possible anchors among the four datasets. Next, “IntegrateData” was used to merge the four individual datasets. After the integration, default clustering steps embedded in Seurat were performed with 20 principal components (PCA) and 3,000 variable genes. The steps included scaling normalized UMI counts with “ScaleData,” dimensional reduction with “RunPCA,” building k-nearest neighbor graph with “FindNeighbors,” and finding clusters with Louvain algorithm by “FindClusters.” Finally, we visualized identified clusters with 2D Uniform Manifold Approximation and Projection (UMAP) by “RunUMAP.” To find the most conserved markers in every cluster, we used “FindConservedMarkers” and show the top 50 markers in **Figure 1F and Table 1**. To evaluate similarities between identified single-cell clusters, we applied unsupervised hierarchical clustering with the pairwise Pearson correlation using 3,000 variable genes (**Figure S1B**). To find DEGs between clusters or conditions, we used “FindMarkers” with padj < 0.05 and log2FoldChange > 0.5 as the thresholds.

### Gene Ontology (GO) Analysis

GO analysis was conducted with DAVID (Huang da et al., 2009; Huang et al., 2009). We used *P* value =0.05 as the threshold to find enriched GO terms such as biological processes.

### Protein–Protein Interaction (PPI) Analysis

A PPI network was drawn with the R package igraph based on the list of reported 1002 PPIs involved in pain (Jamieson et al., 2014). The genes in the DEG lists that had no connections (receive or send) in the PPI network were filtered out. Removed nodes also filtered out any of their edges.

## Results

### Enrichment of DRG Neurons from Pirt-EGFPf Mice for scRNA-seq

Mice were randomly assigned to four groups (datasets): Male-CCI, Female-CCI, Male-Sham, and Female-Sham. Bilateral L4-5 DRGs were collected from mice on day 7 after bilateral sciatic CCI or sham surgery for scRNAseq (**Figure 1A**, **Figure S1A**). In an animal behavior study conducted on day 6 after CCI, paw withdrawal frequencies to low-force (0.07 g) and high-force (0.4 g) mechanical stimulation at the hind paws (data averaged from both sides) were significantly increased (n=5/sex), as compared to the pre-injury frequency, indicating the development of mechanical hypersensitivity (**Figure 1B**). Paw withdrawal frequencies were not significantly changed after sham surgery (n=5/sex).

DRGs contain a large number of non-neuronal cells, including satellite glial cells (SGCs), Schwann cells, immune cells, and fibroblasts. Previous studies have shown that scRNA-seq is advantageous for differentiating neurons from these non-neuronal cells (Renthal et al., 2020; Wang et al., 2021). Here, by using *Pirt*-*EGFPf* mice, we established a highly efficient purification process to further enrich DRG neurons for scRNA-seq (**Figure S1A**).

After removing low-quality cells and doublets, we recovered 3,394 cells from the Male-Sham dataset; 5,678 cells from the Female-Sham dataset; 2,899 cells from the Male-CCI dataset; and 3,681 cells from the Female-CCI dataset. We then utilized canonical correlation analysis embedded in Seurat 3.0 (1), a computational approach for minimizing experimental batch effect, to integrate cells from the four datasets for an unbiased cell clustering. The results showed that the integration worked well in our experiments, as clusters from each dataset aligned well regardless of different biological variations (**Figure 1C**). In total, we identified 16 neuronal clusters and one non-neuronal cluster (**Figure 1D, Figure S1C**).

The non-neuronal cluster had fewer genes and unique molecular identifier (UMI) counts than did neuronal clusters (**Figure S1B)**. It expressed SGC marker genes *Fabp7* and *Apoe* (Renthal et al., 2020; Wang et al., 2021) but showed minimal expression of the pan-neuronal marker *Tubb3*. In contrast, all 16 neuronal clusters showed strong expression of *Tubb3* but had very low levels of *Fabp7* (**Figure S1D**).

Although this non-neuronal cluster showed marker genes known to both Schwann cells and SGCs (e.g., *Mpz*, *Mbp*, *Plp1*), it did not express any Schwann cell-specific genes such as *Mag*, *Prx*, or *Ncmap* (Avraham et al., 2020). Therefore, this non-neuronal cluster included mainly SGCs. No other non-neuronal cluster was detected in our datasets, suggesting a successful enrichment of neurons.

### Prominent New Neuronal Clusters Appear After Sciatic CCI

We further validated the identities of the neuronal clusters by a list of known subtype markers (**Figure 1E**). Our findings confirmed the presence of 12 major standard neuronal clusters [Nppb+ non-peptidergic nociceptors (NP1), Mrgprd+/Cd55+ non-peptidergic nociceptors (NP2), Mrgpra3+/Cd55+ non-peptidergic nociceptors (NP3), Tac1+/Sstr2− peptidergic nociceptors (PEP1-2), Tac1+/Sstr2+ peptidergic nociceptors (PEP3-4), Hpca+/Rarres1+ peptidergic nociceptors (PEP5), Trpv1+/Atp2b4+ peptidergic nociceptors (PEP6), Nefh+/Scn1b+ Aβ low-threshold mechanoreceptors (NF1, NF2), and Fam19a4+/Th+ low-threshold mechano-receptive neurons with C-fibers (cLTMR)]. Importantly, we also identified four new CCI-induced neuronal clusters (CCI-ind1-4) that were prominent in CCI groups but minimal in sham groups (**Figure 1D, E** and **Figure 2A**). CCI-ind1-4 clusters expressed high levels of genes like *Cryba*, *Tgfbi*, *Fgf3*, *Tnfrsf12a*, *Ecel1*, and *Sema6a*, and were clearly segregated from all standard clusters. Expression profiles of the top 50 marker genes in the 16 neuronal clusters are shown in **Figure 1F** and listed in **Table S1**.

**Figure 2.**
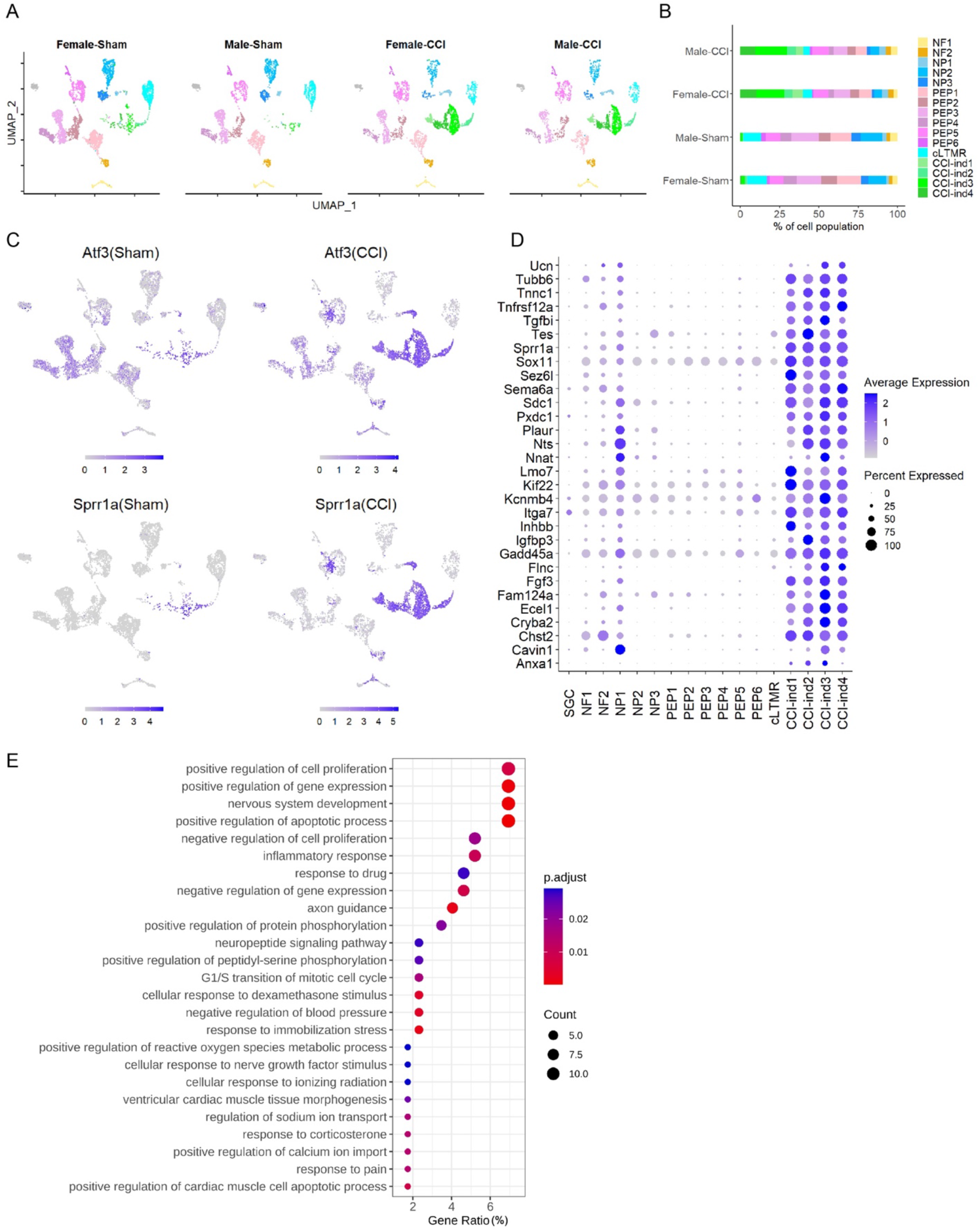
New neuronal clusters induced by CCI of the sciatic nerve. (A) Sciatic CCI induced four new clusters (marked in a red circle) of DRG neurons in both female and male mice. These new clusters were named CCI-ind1, CCI-ind2, CCI-ind3, and CCI-ind4 and were not prominent in sham groups. x-axis: UMAP1, y-axis: UMAP2. (B) Percentage of cell population in 16 neuronal clusters present in each of the four treatment groups. (C) Feature heatmap shows the expression levels of injury-induced genes (*Atf3, Sprr1a*) in different clusters of CCI and sham groups. (D) Dot plot shows the top 30 marker genes of CCI-ind1-4 clusters, as compared to those in other clusters. (E) Top 25 biological processes enriched by the top 50 marker genes from CCI-ind1-4 clusters.

### Gene Programs in CCI-ind1-4 Clusters Are Important to Both Nerve Regeneration and Pain

A much higher percentage of the cell population was contained in the CCI-ind1-4 clusters from CCI groups than from sham groups (**Figure 2A, 2B**). These clusters also showed high expression levels of injury-induced genes such as *Atf3* and *Sprr1a* (**Figure 2C**) (Bonilla et al., 2002; Tsujino et al., 2000). In line with previous findings (Nguyen et al., 2017), *Atf3* was also frequently detected in sequencing of isolated cells from sham or uninjured groups. Comparatively, *Sprr1a* was more selectively expressed in injured neurons of CCI mice (**Figure 2C**). Thus, *Sprr1a* may represent a more specific marker of injured neurons than *Aft3* in scRNA-seq.

Regeneration-associated genes, such as *Anxa1*, *Flnc*, *Gadd45a*, *Inhbb*, *Itga7*, *Kif22*, *Plaur*, *Sema6a*, *Sox11*, *Tnfrsf12a*, and *Tubb6*, were among the top 30 marker genes of CCI-ind1-4 clusters (**Figure 2D**) (Chandran et al., 2016). Intriguingly, some of them may also be involved in neuropathic and inflammatory pain. For example, animal studies suggest that neurotensin (encoded by *Nts*) (Guillemette et al., 2012; Sarret et al., 2005) and annexin1 (encoded by *Anxa1*) may attenuate neuropathic pain and inflammatory pain, respectively (Pei et al., 2011; Zhang et al., 2021). *Nnat* and *Sdc1* were exclusively increased in nociceptive DRG neurons after nerve injury (Chen et al., 2010; Murakami et al., 2015). *Sox11* was identified by integrated bioinformatic analysis as a novel gene that is essential to neuropathic pain (Chen et al., 2021). Moreover, mice that lacked NMDA receptor GluN1 specifically in Schwann cells exhibited upregulation of *Fgf3* and *Sprr1a* in DRG neurons as well as pain hypersensitivity (Brifault et al., 2020); *Fgf3* may also serve as a serum biomarker for sensory neuropathy (Antoine et al., 2015).

Gene ontology (GO) analysis of the top 50 marker genes showed that CCI-induced transcriptomic changes were important for both nerve regeneration (e.g., nervous system development, axon guidance, cellular response to nerve growth factor stimulus) and neuronal excitability (e.g., positive regulation of calcium ion import, response to pain, regulation of sodium ion transport, and the neuropeptide signaling pathway; **Figure 2E**).

### NP1, PEP5, NF1, and NF2 Clusters Exhibit Different Transcriptional Programs From Other Clusters After CCI

Previous studies in different nerve-injury models showed that no specific neuronal cluster was spared from injury. Injured neurons in all clusters lost their original subtype-specific marker genes beginning at day 1 after injury, and hence could no-longer be categorized into original clusters (Hu et al., 2016; Nguyen et al., 2019; Renthal et al., 2020; Wang et al., 2021). Yet, these studies did not further examine the proportion of injured neurons in each cluster after injury. Our findings showed a large decrease of cell proportion in 8 of 12 standard neuronal clusters after CCI (**Figure S2**), suggesting that the injured neurons in these clusters were no longer categorized into their original clusters. Instead, they may be assigned into CCI-ind1-4 clusters (**Figure 3A, B**), as suggested by a previous study (Renthal et al., 2020). The remaining four clusters (NP1, PEP5, NF1, NF2) showed little decrease in cell proportion after CCI. Moreover, a portion of injured neurons (*Sprr1a*^+^) in these four clusters were still categorized into their original clusters after CCI (**Figure 3A, B**). Further analysis revealed two subpopulations of neurons in these clusters (**Figure 3C-F**). For example, in the NP1 cluster, the larger subpopulation showed high expression of *Sprr1a* and other injury-induced genes (*Gal*, *Hspb1*, *Stmn4*, *Chl1*, and *Sox11*) (Chandran et al., 2016), suggesting that these are injured neurons (**Figure 3C**). In contrast, the smaller subpopulation (*Sprr1a*^−^) showed little or no expression of injury-induced genes, but expressed other genes highly, such as *Nppb* and *Sst*, which are NP1 subtype-specific marker genes (**Figure 1E**). We found similar results in PEP5, NF1, and NF2 clusters (**Figure 3D-F**).

**Figure 3.**
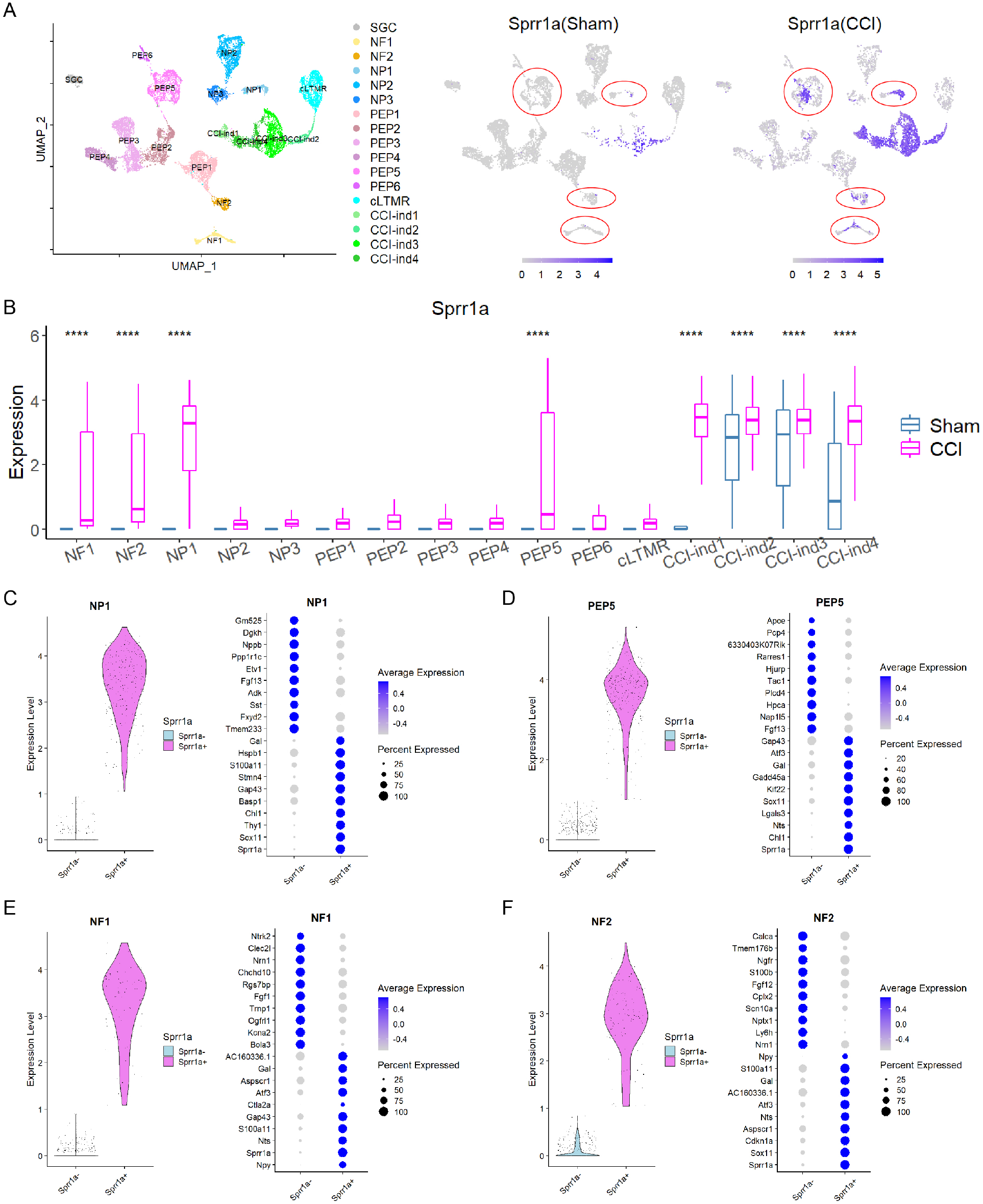
Transcriptional program changes in different neuronal clusters after sciatic nerve CCI. (A) Left: The identities of 17 clusters of DRG cells identified by UMAP. Right: UMAP displays distinct expression patterns of *Sprr1a*, an injury-induced gene, in cells of NP1, PEP5, NF1, and NF2 clusters (indicated by red circles). Each of these clusters contained *Sprr1a*^+^ and *Sprr1a*^−^ cells, resulting in two subpopulations. (B) Box plots show that *Sprr1a* expression was statistically different between sham and CCI groups in only four standard neuronal clusters (NP1, PEP5, NF1, and NF2), suggesting a subtype-specific expression profile in sham and CCI groups. Clusters with average expression level > 1 were selected for statistical analysis. \*\*\*\**P*<0.0001. (C-F) Left: Two subpopulations (*Sprr1a*^+^, *Sprr1a*^−^) of cells in NP1 (C), PEP5 (D), NF1 (E), and NF2 (F) clusters were separated based on the expression of *Sprr1a*. Right: Top 10 DEGs in *Sprr1a*^+^ and *Sprr1a*^−^ subpopulations of NP1, PEP5, NF1, and NF2 clusters.

### Transcriptional Changes in Injured Neurons of NP1, PEP5, NF1, and NF2 Clusters After CCI

Because a significant portion of injured neurons (*Sprr1a*^+^) in NP1, PEP5, NF1, and NF2 clusters maintained their identities after CCI (**Figure 3**), we were able to determine transcriptomic changes in these neurons by comparing them to uninjured neurons of the same clusters in the sham group. After CCI, 197, 67, 41, and 79 differentially expressed genes (DEGs) were generated from NP1, PEP5, NF1, and NF2 clusters, respectively (**Table S2**), with 19 shared DEGs (**Table S3**). Most were common regeneration-associated genes induced by nerve injury (Chandran et al., 2016), and some (*Cacna2d1, Gal, Gap43, Gadd45a, Atf3, Sprr1a*) were also significantly regulated under chronic pain conditions (LaCroix-Fralish et al., 2011; Perkins et al., 2014).

GO analysis showed that these four clusters shared many common pathways, including those related to nervous system development, axon guidance, neuron projection development, microtubule-based process, neuropeptide signaling pathway, and cell differentiation (**Figure 4A**). In addition, we observed CCI-induced changes that affect neuronal excitability (e.g., downregulation of potassium and sodium channels, upregulation of calcium channel Cacna2d1, dysregulation of genes encoding neuropeptide and G-protein-coupled receptors; **Figure S3**).

**Figure 4.**
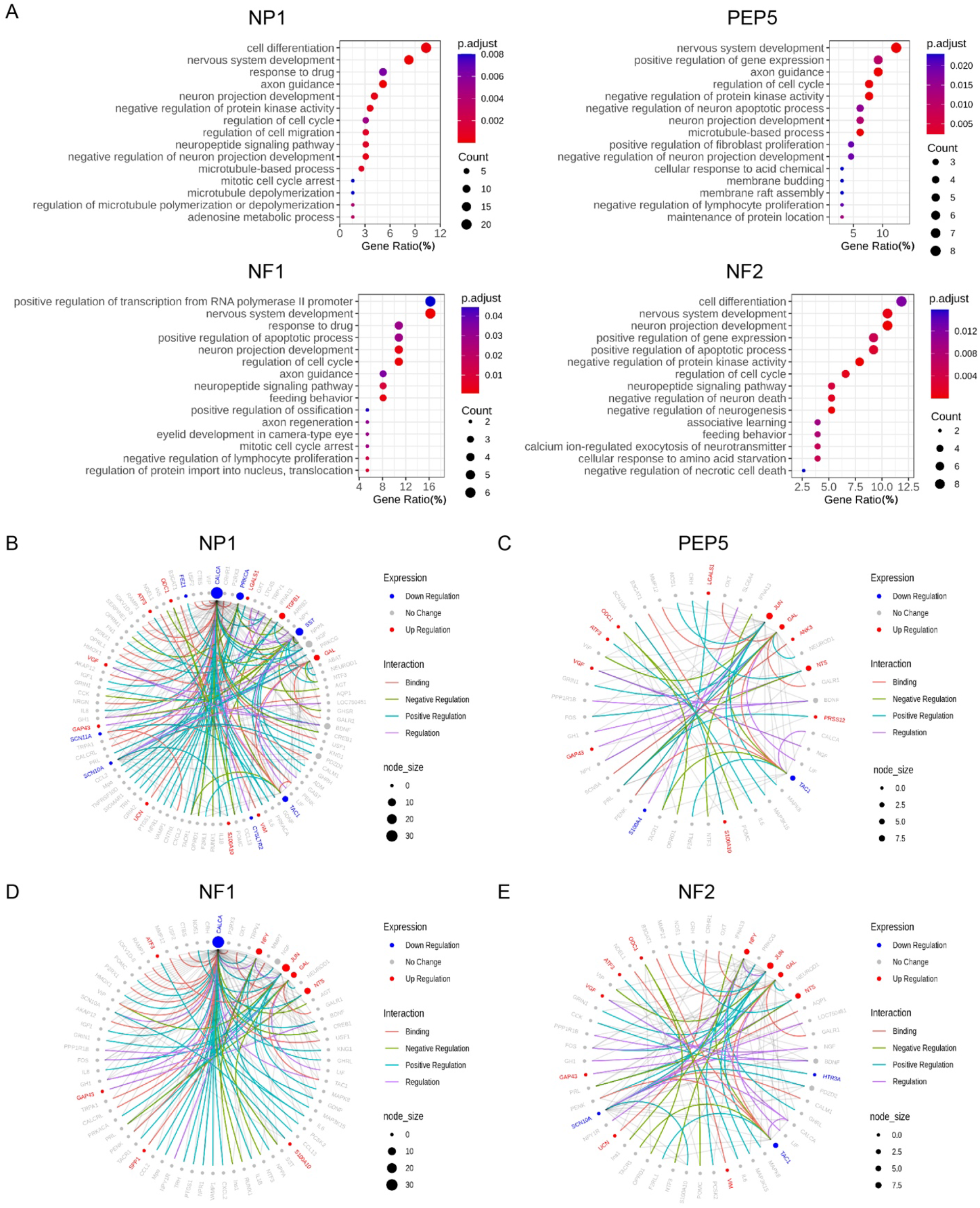
Gene ontology analysis of CCI-induced DEGs and pain-related protein-protein interaction (PPI) networks in NP1, PEP5, NF1, and NF2 clusters. (A) Gene ontology analysis of biological processes enriched by CCI-induced DEGs in NP1, PEP5, NF1, and NF2 clusters. (B-E) The neuropathic pain-specific PPI networks of CCI-induced DEGs in NP1, PEP5, NF1, and NF2 clusters. Colored edges mark the type of interaction. Colored nodes mark the expression changes after CCI. Node size indicates the number of interactions against pain interactome.

We further examined pain-related protein-protein interaction networks within the pain interactome, a comprehensive network of 611 interconnected proteins specifically associated with pain (Jamieson et al., 2014). Examining 197 DEGs of the NP1 cluster revealed an interconnected network of 93 genes (**Figure 4B**). Among them, *Calca, Prkca, Sst*, and *Tac1* were hub genes that were significantly downregulated after CCI, whereas *Tgfb1* and *Gal* were hub genes that were significantly upregulated. Intriguingly, the top marker gene of the NP1 cluster, *Sst*, is also a key gene for neuropathic pain (Zhu et al., 2019); hence it may be an important new target for pain modulation.

When we examined an interconnected network of 44 genes from 67 DEGs in the PEP5 cluster, we identified *Jun, Gal, Nts*, and *Tac1* as hub genes that changed significantly after CCI (**Figure 4C**). Examination of 41 DEGs from the NF1 cluster revealed an interconnected network of 69 genes (**Figure 4D**), with *Calca*, *Npy*, *Jun*, *Gal*, and *Nts* as hub genes. Similarly, an examination of 79 DEGs from the NF2 cluster revealed an interconnected network of 53 genes (**Figure 4E**), including hub genes *Npy*, *Jun*, *Gal*, *Nts*, and *Tac1*.

Among these hub genes, *Calca* and *Tac1* were downregulated whereas *Npy, Gal, Jun*, and *Nts* were upregulated in multiple clusters. Functionally, *Npy* was recently identified as a key prognostic and therapeutic target of neuropathic pain (Tang et al., 2020), and *Jun* and *Nts* may play pivotal modulatory roles in both neuropathic pain and nerve regeneration (Zhao et al., 2020). For example, intrathecal administration of neurotensin (encoded by *Nts*) induced pain inhibition in animal models of neuropathic pain (Guillemette et al., 2012; Sarret et al., 2005). It is possible that the upregulation of *Nts* may represent a compensatory change after injury to limit the exaggeration of neuropathic pain. Future studies are warranted to delineate the roles of each of these shared hub genes in neuropathic pain pathogenesis.

### A Subset of Neuronal Clusters Shows Subtype-Specific Transcriptional Changes in Uninjured Neurons

Another goal of our study was to explore transcriptional changes in uninjured neurons (*Sprr1a*^−^) of different clusters under neuropathic pain conditions. Strikingly, many DEGs were identified in *Sprr1a*^−^ neurons from a subset of clusters including NP1 (143), PEP5 (232), NF1 (95), and NF2 (132) after CCI (**Figure 5A, Table S4**). Comparatively, only a few DEGs were present in *Sprr1a*^−^ neurons from other clusters, suggesting subtype-specific transcriptional changes in uninjured neurons.

**Figure 5.**
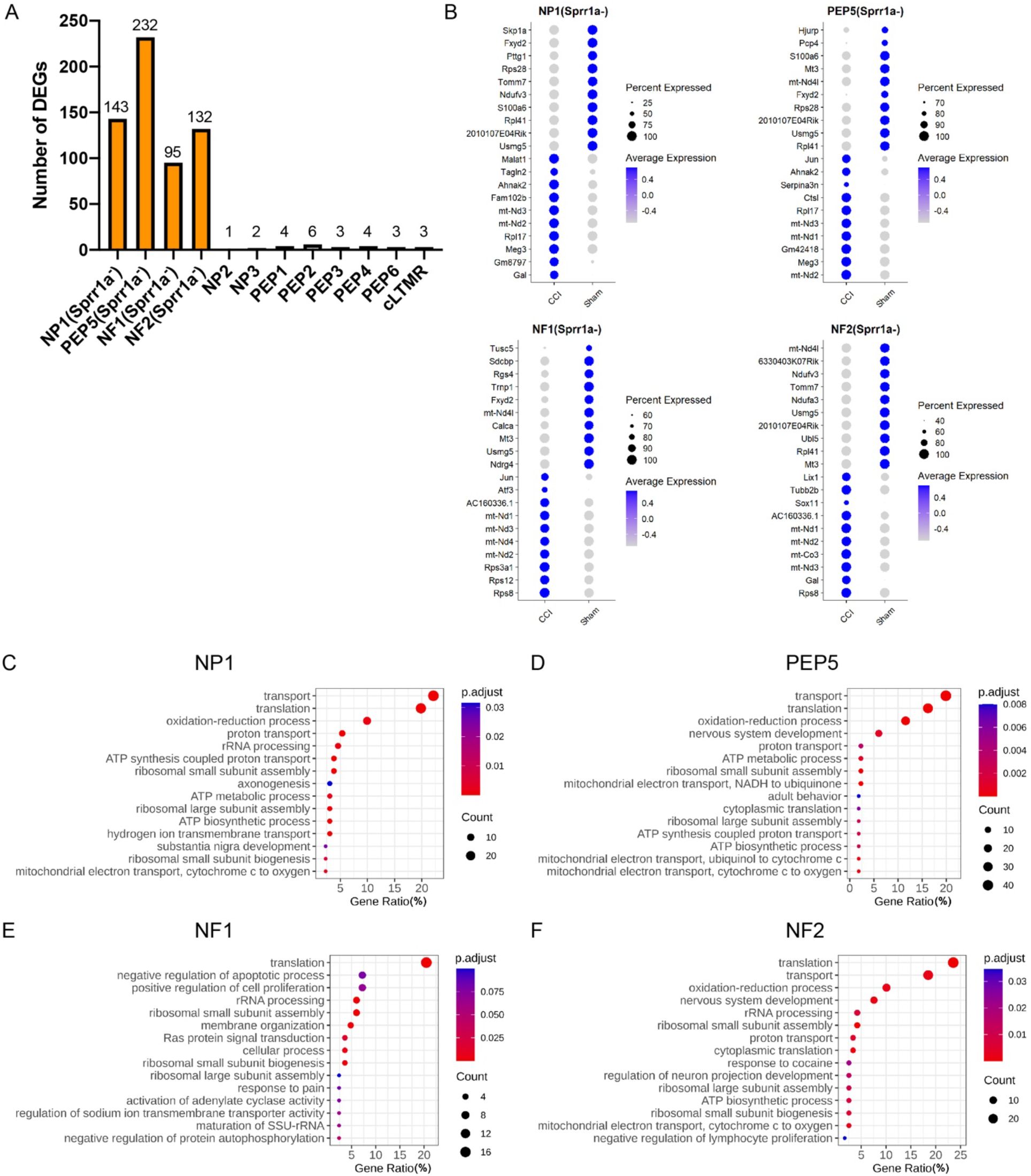
Gene ontology analysis of CCI-induced DEGs in the *Sprr1a*^−^ subpopulation of NP1, PEP5, NF1, and NF2 clusters. (A) Bar graph shows the number of DEGs induced by CCI in *Sprr1a*^−^ neurons of each cluster. (B) Top 10 marker genes of *Sprr1a*^−^ neurons in NP1, PEP5, NF1, and NF2 clusters of CCI and sham groups. (C-F) Gene ontology analysis of CCI-induced DEGs in *Sprr1a*^−^ neurons in NP1 (C), PEP5 (D), NF1 (E), and NF2 (F) clusters.

GO analysis showed that top pathways shared by NP1, PEP5, NF1, and NF2 clusters are related to protein biosynthetic processes, including translation, proton transport, ribosomal small subunit assembly, and transport (**Figure 5B-F, Table S5**). Pathways related to nerve regeneration and neuropathic pain were also found in these clusters. For example, pathways were enriched for genes involved in axonogenesis, ion transport, and mitochondrion transport along microtubule in the NP1 cluster, and for genes involved in nervous system development and regulation of neuron projection development in NF2 and PEP5 clusters.

### Sex Differences in Transcriptional Changes of Different DRG Neuronal Subtypes After CCI

Both human and animal models suggest the presence of sex differences in pain sensitivity and chronic pain prevalence. The peripheral neuronal mechanisms underlying these sexual dimorphisms remain unclear, and few studies have compared transcriptional changes of DRG neurons at the single-cell level, especially under neuropathic pain conditions. *X inactive-specific transcript* (*Xist*) is a specific transcript expressed exclusively by the inactive X chromosome in female mammals (Borsani et al., 1991). Consistent with previous findings, we did not detect *Xist* expression in cells from our male mice (**Figure S4A**). Cell subtype distributions were similar between female and male mice in both sham and CCI groups (**Figure 2B**). Furthermore, Female-Sham and Male-Sham groups showed good correlation in cluster comparison, indicating a great similarity of transcriptional programs under physiologic conditions (**Figure S4B**). Nevertheless, our findings suggest sex differences in the transcriptional changes that occurred after CCI. The Male-CCI group exhibited 303 DEGs when compared to the sham group, and the Female-CCI group expressed 296, with 79 female-specific and 86 male-specific DEGs (**Figure 6A**, **Table S6**). Top pathways were further identified by conducting GO analysis of sex-specific and shared DEGs (**Figure 6B-D**).

**Figure 6.**
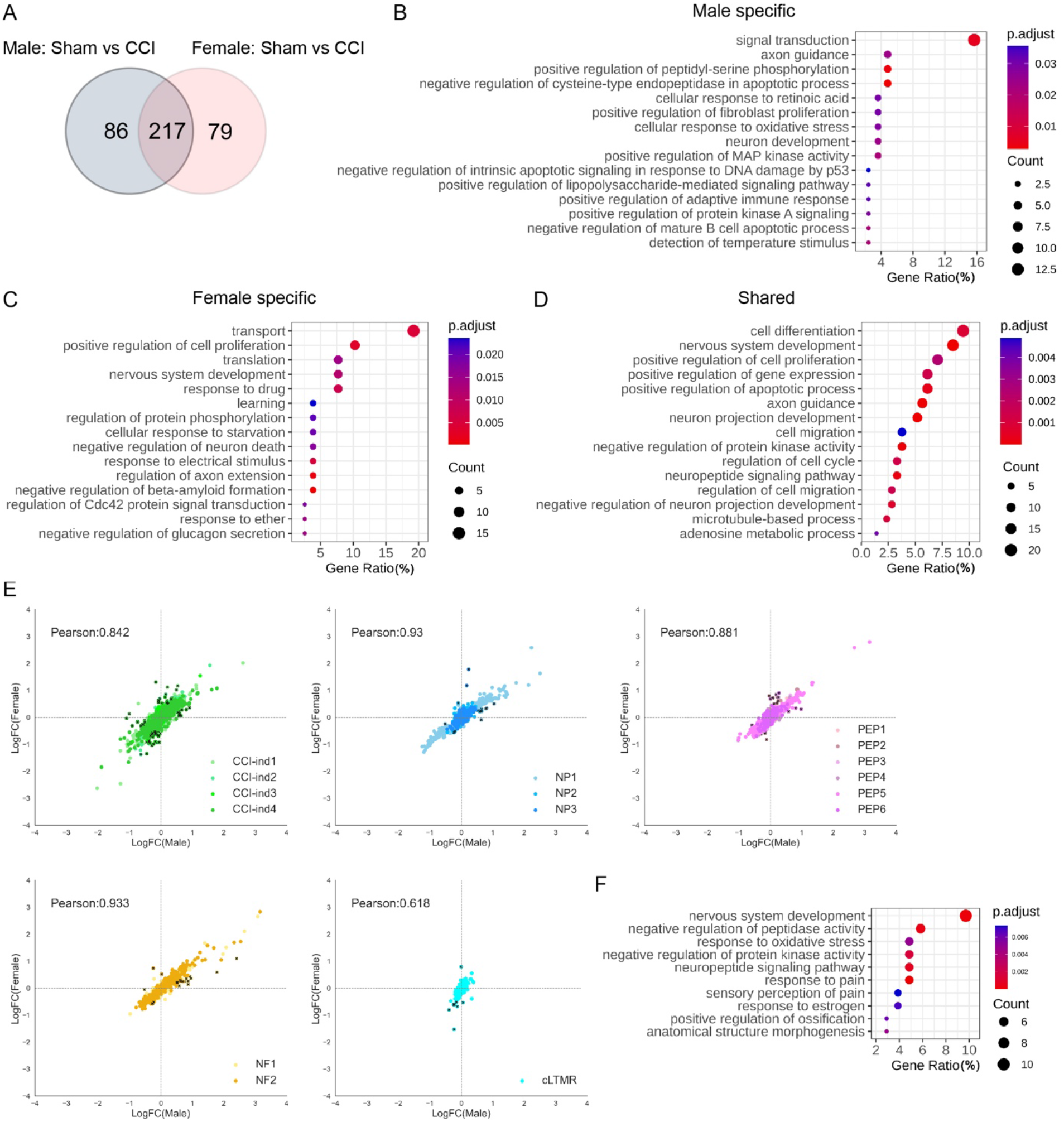
Comparisons of transcriptional changes between female and male mice after CCI. **(A)** Venn diagram shows the number of genes that were differentially expressed between CCI and sham in male and female mice. (B-D) Gene ontology pathways associated with DEGs in male mice only (B), in female mice only (C), and in both male and female mice (D). (E) Pearson correlations based on the fold-change of 382 DEGs after CCI in CCI-ind clusters, NP, PEP, NF, and cLTMR. Black dots represent 106 genes that showed >2-fold differences between female and male mice. (F) Gene ontology analysis of the 106 genes from panel E.

Pearson correlation analysis was also performed based on fold-change of the combined 382 DEGs. Most neuronal clusters showed a good correlation of DEGs between males and females (**Figure 6E**), indicating similarity and a minimal batch effect. Yet, the cLTMR cluster had a poor correlation. Intriguingly, Bohic et al. reported that deletion of *bhlha9*, a transcription factor which is highly expressed in cLTMR, impaired thermotaxis behavior and exacerbated formalin-evoked pain only in male mice (Bohic et al., 2020). Furthermore, *Calca*, which is a nociceptor-specific gene that is highly upregulated in cLTMRs of male *bhlha9*-null mice (Bohic et al., 2020), showed >2-fold differences between female and male cLTMRs in our dataset. These findings suggest that cLTMRs may play an important role in the sexual dimorphism of pain. From all clusters, 106 genes showed >2-fold difference between female and male CCI mice (**Figure 6E, Table S7**). The top 10 pathways from the GO analysis of these DEGs included nervous system development, neuropeptide signaling pathway, response to pain, sensory perception of pain, and response to estrogen (**Figure 6F**). Collectively, these findings suggest differential transcriptomic changes in DRG neurons after CCI between the two sexes.

## Discussion

Primary sensory neurons are the fundamental units of the peripheral sensory system and important for subtype-specific pharmacologic treatment of pain and sensory nerve regeneration. Here, we demonstrated subtype-specific transcriptomic changes in injured DRG neurons under neuropathic pain conditions. In addition to 12 standard clusters validated by known neuronal subtype marker genes, four CCI-ind clusters devoid of subtype marker genes showed a strong presence after CCI. Moreover, we showed that a subset of clusters (NP1, PEP5, NF1, NF2) contain both uninjured (*Sprr1a*^−^) and injured (*Sprr1a*^+^) subpopulations after CCI, but the neurons that retained their subtype marker genes in other clusters were predominantly uninjured. Importantly, we also identified subtype-specific transcriptomic perturbations in uninjured neurons of NP1, PEP5, NF1, and NF2 clusters after CCI. Lastly, our gene expression program uncovered sex differences in the transcriptional changes of DRG neurons after CCI.

Genome-wide screening on bulk DRG tissues has demonstrated profound transcriptional changes after nerve injury (Chandran et al., 2016; LaCroix-Fralish et al., 2011). Yet, because bulk DRG tissue includes a mixture of different neuronal subtypes and non-neuronal cells, RNA-seq cannot distinguish differential transcriptional changes that occur in specific cell subtypes. Recently, scRNA-seq and snRNA-seq studies have begun to uncover subtype-specific perturbations of gene expression in DRG after nerve injury (Nguyen et al., 2019; Renthal et al., 2020; Wang et al., 2021). However, most previous studies isolated cells or nuclei without effectively enriching neurons for sequencing. Consequently, a large number of non-neuronal cells would undergo sequencing reads, thereby reducing sequencing depth (Renthal et al., 2020; Wang et al., 2021). By using *Pirt*-*EGFPf* mice in which EGFP is selectively expressed in most DRG neurons, we improved the purification and successfully enriched DRG neurons for scRNA-seq, with only a few residual SGCs; most other non-neuronal clusters were excluded. The resulting increase in sequencing depth increased the number of DEGs detected in DRG neurons, particularly for genes with low read counts (Perkins et al., 2014).

Clustering analysis identified 16 distinct neuronal clusters and one SGC cluster. Of those, 12 standard neuronal clusters were categorized based on known subtype marker genes, which were present in both sham and CCI groups. These findings suggest that a subpopulation of neurons in each subtype was spared from injury and maintained its distinguishing transcriptional program at day 7 post-CCI, when neuropathic pain has reached a peak in this model (Bennett & Xie, 1988). Strikingly, CCI-ind1-4 clusters showed diminished expression of subtype marker genes but high expression of injury-induced genes (*Atf3*, *Sprr1a*). These clusters were present at high levels only after CCI. Accordingly, they are likely injured neurons that have lost their original subtype marker genes. Indeed, the top 50 DEGs in CCI-ind1-4 clusters included those important to nerve regeneration and neuronal hyperexcitability. Because these neurons are likely axotomized and disrupted from peripheral receptive fields, they may play important roles in nerve regeneration and spontaneous pain after nerve injury. These findings are consistent with previous results obtained with snRNA-seq, which showed a new transcriptional state in DRG neurons after spared nerve injury and in trigeminal ganglion neurons after infraorbital nerve transection (Nguyen et al., 2019). A similar phenomenon was noted after spinal nerve transection, sciatic nerve transection, and crush injury (Renthal et al., 2020; Wang et al., 2021). The common characteristics of these new neuronal clusters are the increased expression of injury-induced genes and diminished original neuron subtype-specific marker genes such as *Tac1*, *Mrgprd*, *Nefh*, *Th*, and *Sst*, which may represent a general adaptation mechanism after mechanical/traumatic nerve injuries. The activation of new transcriptional programs in these neurons may help shift from normal functions (e.g., neurotransmission) to protection and regeneration modes.

Unlike *Atf3*, which was frequently detected in isolated cells even in the sham group, *Sprr1a* showed high specificity as an injury indicator in our scRNA-seq. Even though *Sprr1a* was used to differentiate injured and uninjured neurons, a small number of *Sprr1a*^+^ neurons and CCI-ind1-4 clusters also appeared in sham groups. Their presence may have been induced by the skin incision and muscle damage of sham surgery, which can produce minor nerve injury, or by the cell stress and injury that occur during tissue harvesting and processing (e.g., dissociation, enzymatic digestion). A previous study showed that even sham surgery and minor peripheral injury may increase *Atf3, Sox11, Sema6a, Csf1*, and *Gal* in DRG neurons (Nguyen et al., 2019).

Another salient finding is that a portion of injured neurons (*Sprr1a*^+^) in NP1, PEP5, NF1, and NF2 clusters maintained their original neuronal identities after CCI, and hence can still be clustered together with uninjured neurons. In contrast, the other eight clusters contained only uninjured neurons after CCI. Thus, a subpopulation of injured neurons in these four clusters may exhibit different transcriptomic changes after CCI than injured neurons in other clusters. The injured neurons in other clusters may have lost their identities, entered a new transcriptional state, and been assigned to CCI-ind1-4 clusters. This notion is supported by decreased cell populations in these clusters after CCI and a recent snRNA-seq study that reclassified subtypes of injured neurons by examining multiple post-injury time points to consecutively capture residual transcriptional signatures (i.e., a set of processed transcripts that are expressed in a specific cluster) during the transition from uninjured to injured states (Renthal et al., 2020). It remains to be determined why injured neurons in NP1, PEP5, NF1, and NF2 clusters can still be clustered together with uninjured neurons from the same subtype, even though they have also lost some subtype-specific marker genes. This apparent discrepancy may be due to these neurons losing expression of only a subset of marker genes after CCI, but maintaining the major transcriptional signatures of the naïve state, which distinguishes them from other neuronal clusters.

Increasing evidence has suggested that uninjured DRG neurons also play important roles in neuropathic pain, and show robust neurochemical and functional changes after nerve injury (Kalpachidou et al., 2021; Obata et al., 2003; Pertin et al., 2005; Tran & Crawford, 2020). Strikingly, our analysis showed for the first time that uninjured (*Sprr1a*^−^) neurons in NP1, PEP5, NF1, and NF2 clusters also undergo subtype-specific changes in gene expression after CCI, as compared to those in the sham group. GO analysis suggested that these clusters underwent common changes in pathways involved in protein biosynthesis (e.g., translation, ribosomal small subunit assembly) and differential changes in pathways related to regeneration, pain, axonogenesis, and ion transport. Functional changes of uninjured neurons under neuropathic conditions may extend beyond transcriptional regulation to also involve mechanisms such as increased translational rate (Gebauer & Hentze, 2004) and modulations by miRNA. In line with this notion, our GO analysis showed that a pathway named “*positive regulation of Pri-miRNA transcription from RNA polymerase promoter*” was significantly regulated in the NF2 cluster. Systematic investigations are needed to sort out the functional effects of transcriptomic changes in injured and uninjured neurons from each cluster on nerve regeneration, neuronal excitability, and pain. We focused on examining transcriptional changes at day 7 post-CCI, when neuropathic pain-like behavior in mice reaches a peak and enters the maintenance phase. Since gene expression changes may vary at different neuropathic pain stages, a time course study of transcriptional changes in different subtypes of DRG neurons after CCI is warranted.

*Xist* is known to be expressed exclusively by the inactive X chromosome in mammals (Borsani et al., 1991). Indeed, we also found that it was selectively expressed in female but not male datasets, suggesting that *Xist* is a reliable female marker gene for scRNA-seq. Pearson correlation analysis suggested a similarity of gene expression program in the two sexes under physiologic conditions. However, we found sex differences in transcriptional changes after CCI, and the cLTMR cluster may play an important role in the sexually dimorphic pain response. Among 296 female and 303 male DEGs generated after CCI, 79 were female-specific and 86 were male-specific. In addition, 106 genes showed more than 2-fold differences between the two sexes after CCI. These findings are consistent with a bulk RNA-seq study of DRG, which showed vast differences in genes enriched in pain-relevant pathways between female and male rats after CCI (Stephens et al., 2019). Although these findings suggest that peripheral neuronal mechanisms may also underlie sexual dimorphisms in neuropathic pain, Renthal et al. reported no differences in subtype distributions or injury-induced transcriptional changes between males and females after sciatic nerve crush injury (Renthal et al., 2020). The reasons for this discrepancy are unclear. It may be partially due to differences in the techniques (e.g., tissue processing, cell sorting, sequencing) and animal models. The read count and sequencing depth were lower in the previous study, which may have resulted in fewer DEGs and decreased ability to detect subtle changes. Details of the distinct gene networks that function in different neuronal subtypes and might underpin sexual dimorphisms in neuropathic pain must still be explored at a single-cell level.

## Conclusions

In summary, our findings in a well-established animal model of neuropathic pain share some similarities with recent findings in transection injury models, including the loss of marker genes in injured neurons and the emergence of new, injury-induced clusters. Importantly, we demonstrated subtype-specific transcriptomic changes in both injured and uninjured neurons of NP1, PEP5, NF1, and NF2 clusters after CCI. Furthermore, transcriptomic sexual dimorphism may occur in DRG neurons after nerve injury, and cLTMR may play a pivotal role in sex-specific pain modulation. This new knowledge may provide important rationales for developing cell subtype-specific and sex-based therapies that can optimize neuropathic pain inhibition and sensory nerve regeneration. An ideal therapy would target uninjured neurons in NP1, PEP5, NF1, and NF2 clusters for pain control but would not interfere with important transcription programs needed for regeneration of injured neurons.

## Acknowledgements

This study was conducted at the Johns Hopkins University School of Medicine. The authors thank Claire F. Levine, MS (scientific editor, Department of Anesthesiology and Critical Care Medicine, Johns Hopkins University), for editing the manuscript. This study was supported by National Institutes of Health (Bethesda, Maryland, USA) grants NS070814 (Y.G.), NS110598 (Y.G.), and NS117761 (Y.G.) and was facilitated by the Pain Research Core, which is funded by the Blaustein Fund and the Neurosurgery Pain Research Institute at the Johns Hopkins School of Medicine. The authors thank Dr. Xinzhong Dong for sharing the *Pirt*-*EGFPf* mouse strain. The authors thank Hao Zhang from the Flow Cytometry Cell Sorting Core Facility at Bloomberg School of Public Health, Johns Hopkins University for doing FACS sorting. The facility was supported by 1S10OD016315-01,1S10RR13777001 and in part by CFAR: 5P30AI094189-04 (Chaisson). The authors appreciate the Johns Hopkins Transcriptomics and Deep Sequencing Core for conducting the single-cell RNA-seq. Funders had no role in study design, data collection, or data interpretation, or in the decision to submit the work for publication. The authors declare no competing interests. There are no other relationships that might lead to a conflict of interest in the current study.

## Supplemental figures and legends

**Figure S1.**
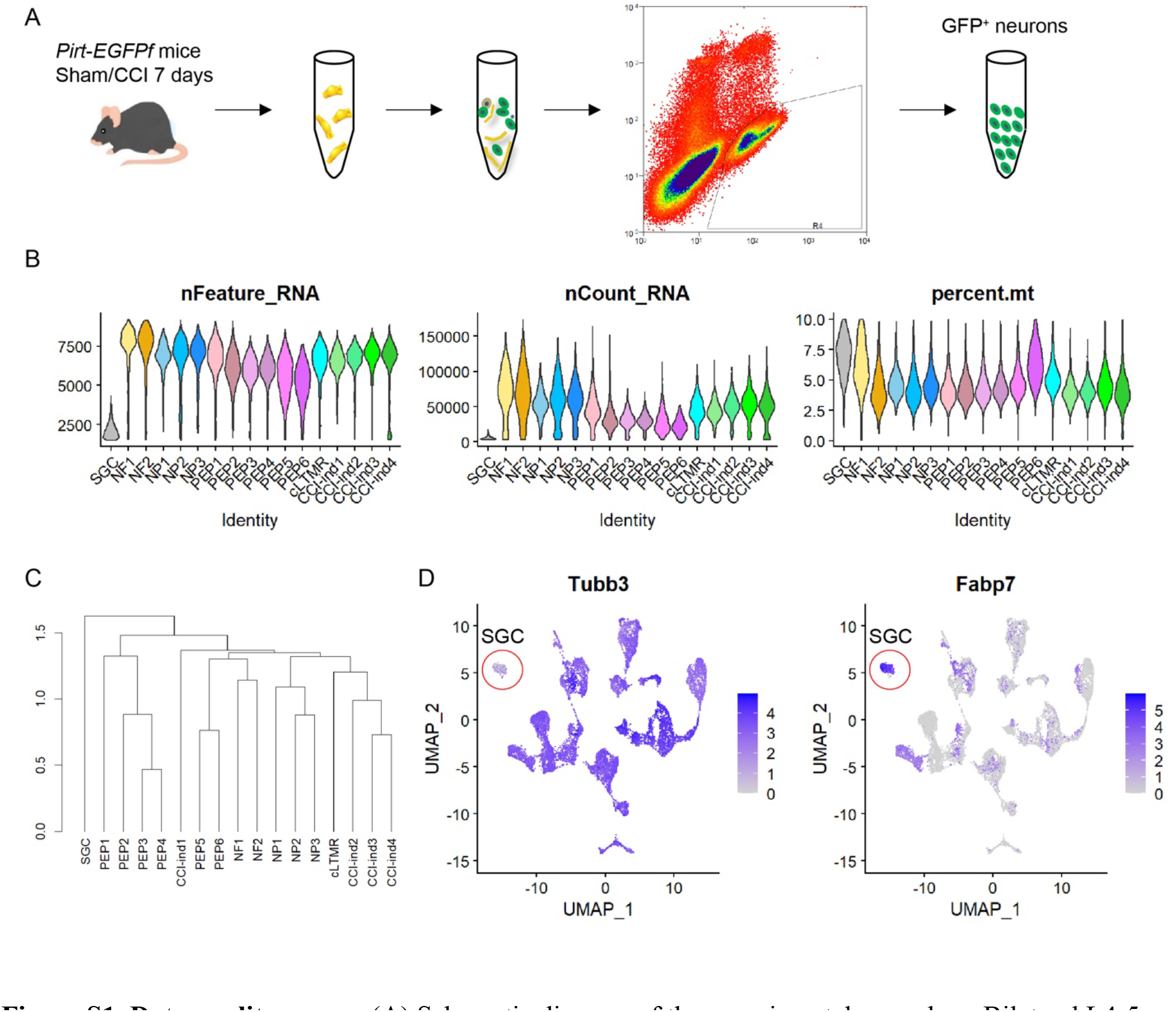
Data quality assays. (A) Schematic diagram of the experimental procedure. Bilateral L4-5 DRGs were dissected from *Pirt*-*EGFPf* mice on day 7 after sham surgery (sham) or bilateral chronic constriction injury (CCI) of the sciatic nerve. The dissociated cell suspension was processed with flow cytometry to collect GFP^+^ cells. (B) Violin plots show the number of expressed genes (left), UMI counts (middle), and percent of mitochondrial genes in each cell cluster (right). (C) Hierarchical clustering of 17 cell clusters identified in the DRG. (D) Feature heatmaps show the expression of *Tubb3* (a pan-neuronal marker) and *Fabp7* [a satellite glial cell (SGC) marker] in all clusters. The SGC cluster is indicated with a red circle. The color scale indicates the Log2 normalized transcript counts.

**Figure S2.**
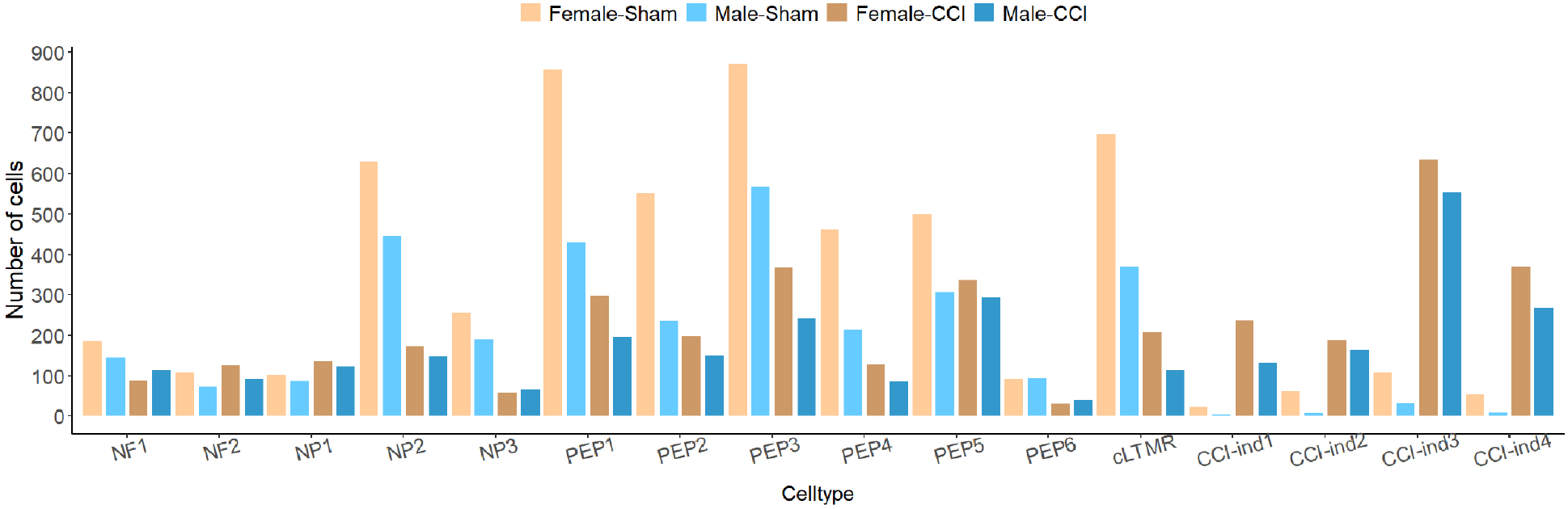
Cell number of each neuronal cluster in the four datasets.

**Figure S3.**
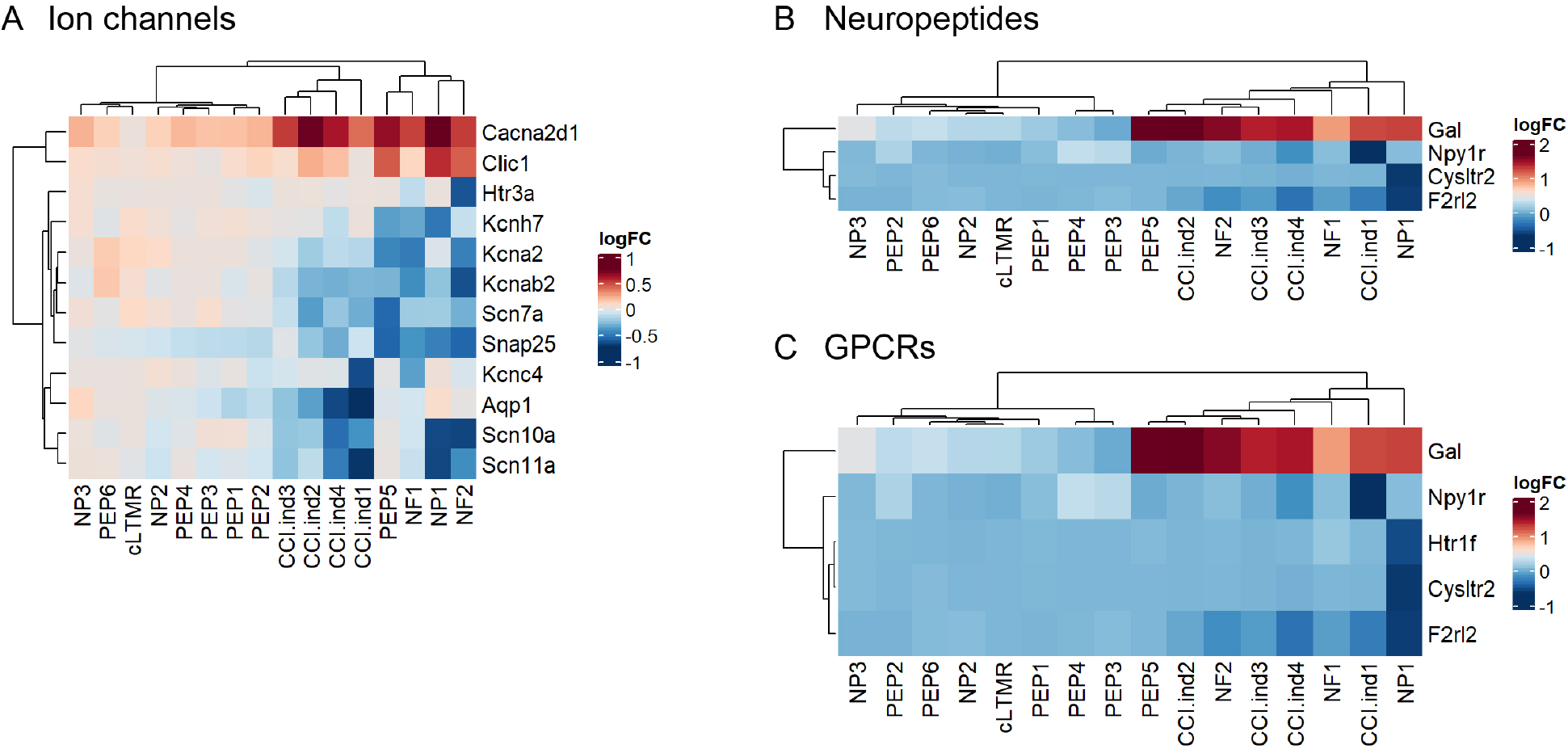
CCI altered the expression of genes that encode ion channels, neuropeptides, and G-protein-coupled receptors (GPCRs) in DRG neurons. (A-C) Heatmaps of the log2FC (CCI neurons compared to sham neurons for each cell type) of select genes encoding ion channels (A), neuropeptides (B), and G-protein-coupled receptors (GPCRs) (C). Genes shown on the heatmap are significantly regulated after CCI.

**Figure S4.**
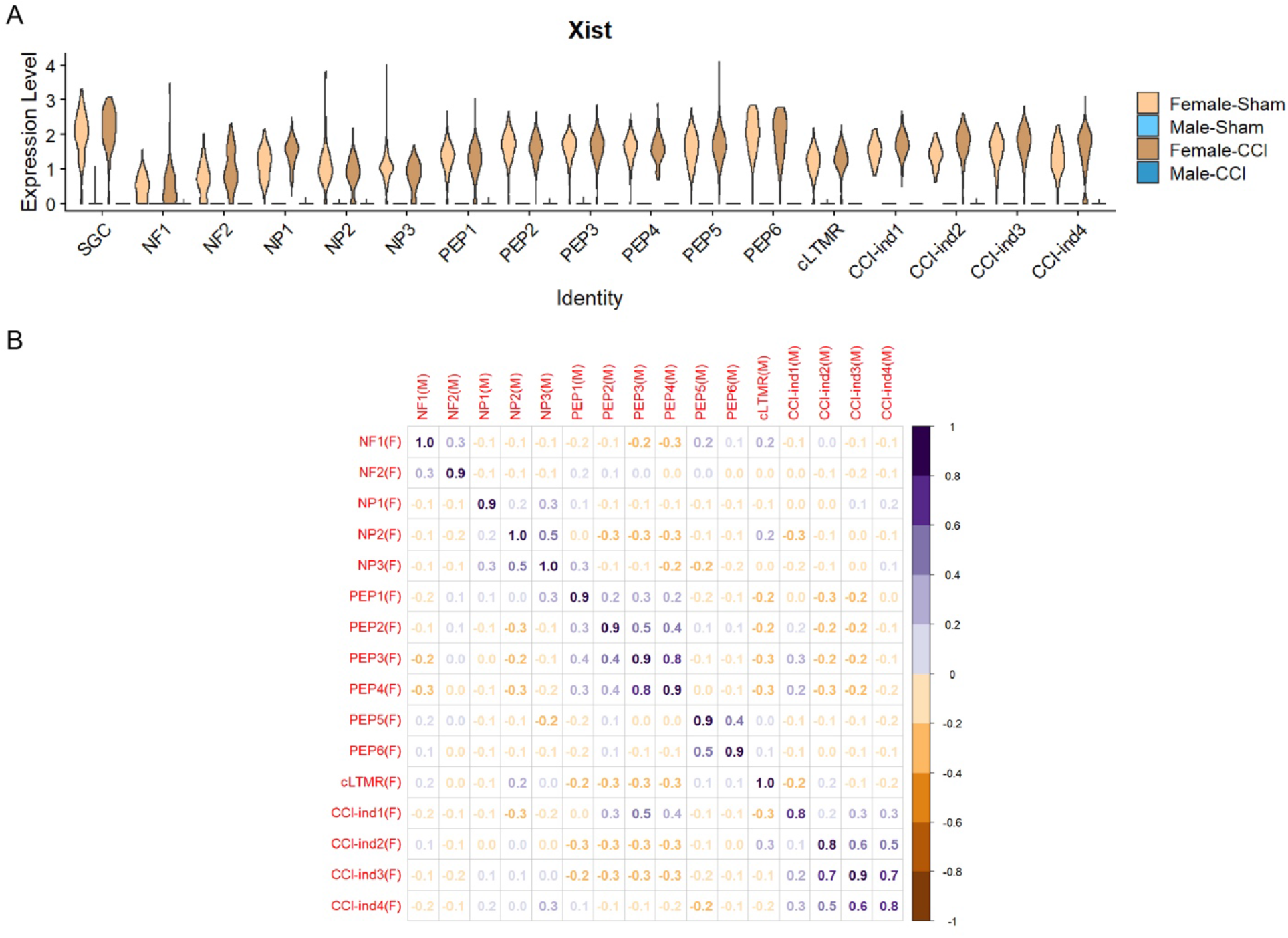
Comparisons of specific DEGs between female and male mice after CCI. (A) Violin plot shows the expression levels of *Xist* in each cell cluster in the four groups/datasets. (B) Pearson correlation of each neuronal cluster between Male-Sham and Female-Sham mice, based on whole transcript counts.

**Supplementary Table 1 Dataset**

Top 50 conserved marker genes in 16 neuronal clusters (related to Figure 1).

**Supplementary Table 2 Dataset**

Differentially expressed genes in NP1, PEP5, NF1, NF2, comparing CCI with Sham (padj<0.05) (related to Figure 4)

**Supplementary Table 3 Dataset**

19 differentially expressed genes shared by NP1, PEP5, NF1 and NF2, comparing CCI with Sham (related to Figure 4).

**Supplementary Table 4 Dataset**

Differentially expressed genes in Sprr1a^−^ neurons of NP1, PEP5, NF1 and NF2, comparing CCI with Sham (padj<0.05) (related to Figure 5).

**Supplementary Table 5 Dataset**

Enriched pathways after CCI in NP1, PEP5, NF1 and NF2 with the corresponding genes (related to Figure 5).

**Supplementary Table 6 Dataset**

Lists of CCI-induced differentially expressed genes including DEGs only in male, DEGs only in female and DEGs shared by both male and female (padj<0.05, Log2Fold change>0.5) (related to Figure 6).

**Supplementary Table 7 Dataset**

Enriched pathways with the 106 CCI-induced DEGs (related to Figure 6).

